# Strategies for identifying dynamic regions in protein complexes: flexibility changes accompany methylation in chemotaxis receptor signaling states

**DOI:** 10.1101/2020.03.03.974428

**Authors:** Nikita Malik, Katherine A Wahlbeck, Lynmarie K Thompson

## Abstract

Bacterial chemoreceptors are organized in arrays composed of helical receptors arranged as trimers of dimers, coupled to a histidine kinase CheA and a coupling protein CheW. Ligand binding to the external domain inhibits the kinase activity, leading to a change in the swimming behavior. Adaptation to an ongoing stimulus involves reversible methylation and demethylation of specific glutamate residues. However, the exact mechanism of signal propagation through the helical receptor to the histidine kinase remains elusive. Dynamics of the receptor cytoplasmic domain is thought to play an important role in the signal transduction, and current models propose inverse dynamic changes in different regions of the receptor. We hypothesize that the adaptational modification (methylation) controls the dynamics by stabilizing a partially ordered domain, which in turn modulates the binding of the kinase, CheA. We investigated the difference in dynamics between the methylated and unmethylated states of the chemoreceptor using solid-state NMR. The unmethylated receptor (CF4E) shows increased flexibility relative to the methylation mimic (CF4Q). Methylation helix 1 (MH1) has been shown to be flexible in the methylated receptor. Our analysis indicates that in addition to MH1, methylation helix 2 also becomes flexible in the unmethylated receptor. In addition, we have demonstrated that both states of the receptor have a rigid region and segments with intermediate dynamics. The strategies used in the study for identifying dynamic regions are applicable to a broad class of proteins and protein complexes with intrinsic disorder and dynamics spanning multiple timescales.

**Graphical Abstract:** 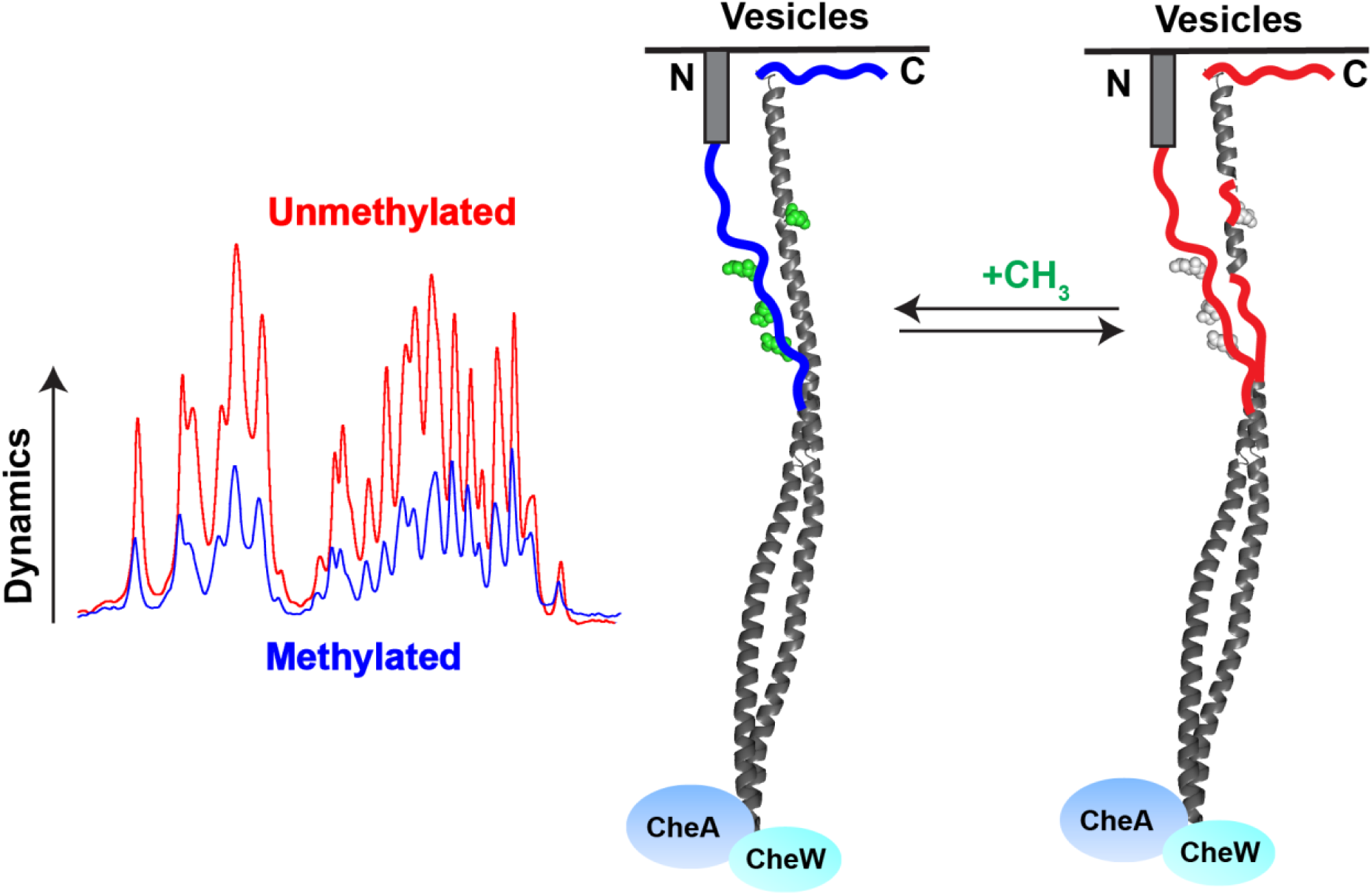

**Highlights:**

- Receptors exhibit greater ns timescale dynamics in unmethylated vs methylated state
- Methylation helix 2 likely involved in increased flexibility of unmethylated state
- Dynamics occur on multiple timescales in both states of the receptor

## Introduction

Most biological processes are dependent on proteins interacting with other proteins and assembling into complexes[1]. Large protein complexes, membrane proteins, and dynamic proteins are challenging targets for structural biology. Complexes involving dynamic or partially disordered proteins can be difficult to study using traditional methods because of quick association and dissociation kinetics due to flexible binding partners and/or conformational rearrangement in complexes[2–4]. The challenge of crystallizing disordered proteins and membrane proteins limits the utility of x-ray crystallography, and the slow tumbling of large protein complexes limits the utility of solution NMR. Cryo-electron microscopy can provide complementary structural information on such systems, but not for domains with extensive dynamics or disorder. Solid-state NMR is ideal for studies of large protein complexes and membrane proteins because it is not reliant on long range order, solubility, or rapid tumbling[5]. In the past decade, solid-state NMR has been used to study a variety of biological systems such as HIV-1 capsid protein assemblies[6–11], 300 kDa GB1-antibody complexes[12,13], and membrane-bound influenza M2 proton channels[14–16]. Such NMR studies have also provided measurements of protein dynamics [17–19] and identification of disordered regions [20–24].

Research in recent years on intrinsically disordered proteins (IDPs) and intrinsically disordered regions (IDRs) has highlighted the importance of disordered regions in the function of many proteins. These regions can be essential for correct function, by mediating protein-protein interactions, protein-DNA interactions, and protein-RNA interactions[25]. They often act as phosphorylation sites in proteins, which in turn act as protein interaction sites[26,27]. Conformational flexibility can allow the protein, for example the transcriptional activator CBP, to bind to both its target and to modifying enzymes[28]. Intrinsic disorder is also an important component of cellular signaling[26], because of the favorable characteristics of IDPs. IDPs have advantages in kinetics because fast association rates allow signals to rapidly turn on[26]. Alternatively, IDPs can bind partners with high specificity and average affinity, enabling the signal to rapidly terminate with dissociation of ligands[26]. Because many proteins have both structured and intrinsically disordered regions, assessing the dynamics and identifying dynamic regions is an important first step in characterizing any protein[26,29,30].

In this study, we investigate the dynamics of a protein involved in cellular signaling, the *E coli* aspartate receptor, which we have recently proposed contains a partially disordered domain involved in the signaling mechanism[31]. Chemotaxis receptors (reviewed in [32,33]) help bacteria sense and respond to their environment. These proteins, embedded in the membrane of the cell, detect chemical gradients in the environment and control activity of a histidine autokinase, CheA. This kinase phosphorylates the response regulator CheY, which interacts with the flagellar motor to alter cell swimming. As these receptors help sense their environment to bias the swimming of the cell towards nutrients and away from toxic molecules, they are important for the survival and propagation of bacteria. Chemotaxis receptors have helical secondary structure and are composed of 3 domains: the ligand-binding periplasmic domain, the HAMP domain, and the cytoplasmic domain. In their native form, the receptor proteins form trimers of dimers that further arrange into hexagonal nanoarrays. The cytoplasmic partners CheA and CheW form rings that connect the receptors at their membrane-distal tips[34–36]. Kinase activity is modulated through two processes: ligand binding and methylation. Sensing a chemical gradient in the cell’s environment begins when the periplasmic end of the receptor binds small molecules[37]. This binding event induces a 2 Å piston motion of an alpha helix that extends through the transmembrane region of the receptor[38]. Changes in receptor dynamics are thought to be involved in propagating the signal an additional 200 Å through the HAMP and cytoplasmic domains of the receptor to modulate the activity of the associated kinase CheA[39,40]. After attractant binding turns off CheA, methylation of glutamic acid residues in the cytoplasmic domain of the receptor by the methyl-transferase CheR restores the kinase-ON state. Demethylation by the methyl-esterase CheB shifts the receptor to the kinase-OFF state Receptor methylation and demethylation, which enables the receptor to adapt to an ongoing stimulus, is an essential component of chemotaxis.

INEPT and CP experiments have been used routinely in solid-state NMR to obtain information about the overall dynamics in a protein[41]. The scalar-based INEPT experiment is used to observe highly flexible regions of the protein that have longer transverse relaxation (T_2_) values. The signal from the rigid portions with shorter ^1^H T_2_ decays during the INEPT delays, leaving observable signal only from the segments with mobility on the ns or shorter timescale. The rigid regions exhibit strong dipole-dipole coupling and are observed by the cross-polarization (CP) experiment. The combination of CP and INEPT experiments has successfully characterized many systems including phospholamban in lipid bilayers[42], amyloid beta fibrils[43], and chicken α-spectrin SH3[44]. Experiments probing dynamics at other timescales include measurement of relaxation parameters like T_1p_ [45,46], and DIPSHIFT or REDOR measurement of motional averaging of anisotropic parameters [46,47].

Results of mutagenesis studies have led to proposals that dynamics likely have a role in the mechanism of chemoreceptor signal transmission[39,40]. Furthermore, we have recently proposed that the receptor cytoplasmic domain is partially disordered and that signal propagation involves modulation of this disorder[31]. Solid-state NMR studies can be used to assess the dynamics of each receptor signaling state and then determine the best NMR approaches for future experiments. In previous studies, we have conducted CP vs. INEPT experiments on the kinase-ON and kinase-OFF state of the cytoplasmic fragment of the receptor in complex with CheA and CheW. We reported that the methylation helix 1 (MH1) and the C-terminal tail have significant dynamics on the ns-timescale or shorter in both signaling states[48]. We have also probed the ms-timescale dynamics of the cytoplasmic fragment in functional complexes using carbon-nitrogen REDOR experiments. There we found that the receptor backbone carbons have only 60%-75% of the expected REDOR dephasing, the other 40-25% of the CP-observable portions of the protein backbone has mobility that averages ^13^C^15^N dipolar couplings to zero, and the extent of this mobility changes with signaling state[47]. We have now compared the dynamics between the unmethylated and methylated states of the chemotaxis receptor cytoplasmic fragment to probe the mechanism of adaptation. CP vs INEPT spectra show increased dynamics in the demethylated state of the receptor, and some of these increases are due to additional region(s) of the receptor becoming highly dynamic. Based on analysis of the residue types and resonance intensities, we propose this region is methylation helix 2 (MH2). In addition, we have shown through measurement of CP efficiency and buildup curves that parts of the receptor have intermediate timescale dynamics. These methods are applicable to other protein systems to identify the presence of flexible or disordered regions, with dynamics on multiple timescales.

## Methods

### Protein Purification

The cytoplasmic fragment (CF) of the *E. coli* aspartate receptor was isotopically labeled and expressed in BL21(DE3) E. coli cells using plasmids encoding CF with an N-terminal His-tag as previously described[49]. The plasmid pHTCF4Q (amp^r^) encodes CF with Gln at all 4 methylation sites, mimicking the methylated state of the receptor, and pHTCF4E (amp^r^) encodes CF with Glu at those sites, representing the unmethylated receptor. The CF expression plasmids do not contain lacI^q^; hence they were co-transformed with pCF430 (tet^r^) encoding lacI^q^. Cells were grown as previously described[48] in M9 medium supplemented with ^13^C-glucose and (^15^NH_4_)_2_SO_4_ as the sole carbon and nitrogen source, respectively. The proteins were purified using Ni-affinity chromatography and the concentrations were estimated using BCA assay as previously described[49].

Plasmids pTEV-cheA (kan^r^) and pTEV-cheW (kan^r^), encoding CheA and CheW respectively, were expressed in BL21(DE3) cells. Each unlabeled protein was purified using Ni-affinity chromatography and the TEV cleavable N-terminal His-tag was removed as previously described[49]. The extinction coefficients of 25,000 M^-1^cm^-1^ for CheA and 5120 M^-1^cm^-1^ for CheW were used to measure the protein concentrations using A_280_[50,51].

### NMR sample preparation

The functional membrane bound complex of the receptor (both CF4Q and CF4E) and the two binding partners CheA and CheW was assembled as previously described. Briefly, the vesicles were composed of a 1:1.5 ratio of the lipids DOGS-NTA-Ni^2+^ (1,2-dioleoyl-sn-glycero-3-[(N-(5-amino-1-carboxypentyl) iminodiacetic acid)succinyl] (nickel salt) and DOPC (1,2-dioleoyl-sn-glycero-3-phosphocholine) (Avanti Polar Lipids); after hydration and five freeze-thaw cycles, vesicles were extruded through a 100 nm pore polycarbonate membrane[49]. Complexes were assembled (4 ml for each NMR sample) by adding 1mM PMSF, 12uM CheA, 24uM CheW, 30uM CF (^13^C,^15^N labeled), and 725uM vesicles to the kinase buffer (50 mM K_x_H_x_PO4, 50 mM KCl, 5 mM MgCl_2_, pH 7.5) and incubated overnight at 25°C, followed by checking the activity and sedimentation[49]. The functional complex was recovered as a jelly-like pellet by centrifugation at 60,000 rpm at 25°C for 2 h in an ultracentrifuge (128,000 g in a TLA 120.2 Beckman rotor) and packed into a 1.9 mm rotor. The assembly conditions yielded reproducible amounts of complexes for CF4Q and CF4E samples; 13-14 mg of pellet was packed in each rotor. The mass of CF in each rotor was calculated as previously described[48] and the mass difference was accounted for during data analysis and signal intensity calculations.

A sacrificial sample for temperature calibration was made by adding unlabeled CF4E and 20 mM TmDOTP to the lipid vesicles prepared in deuterated kinase buffer.

### NMR Spectroscopy

All experiments were recorded on a Bruker Avance III spectrometer (14.1T magnet) with a 1.9mm HNCF MAS probe (^1^H 600 MHz, ^13^C 150 MHz, and ^15^N 60 MHz). For the CF4Q sample at 12 °C and 11.11kHz MAS, π/2 pulse lengths of 3.6 μs for ^1^H and 4.5 μs for ^13^C were used. ^1^H-^13^C cross polarization was performed with ^13^C B1 of 55 kHz, ^1^H B1 of 67 kHz (30% ramp centered close to the +1 condition) for a contact time of 0.9 ms, followed by SPINAL64 decoupling at 70 kHz and a 5 s recycle delay. The parameters for the CF4E sample were similar to CF4Q with the exception of the ^1^H 90° pulse at 3.4 μs, SPINAL64 decoupling at 73 kHz, and the ^1^H B1 during CP at 44 kHz (30% ramp centered on the −1 condition). Experiments on frozen samples used the same parameters except ^1^H π/2 pulse lengths were 3.2 μs (CF4Q) and 3.1 μs (CF4E), and the re-optimized cross polarization was performed with ^1^H B1 of 70 kHz (CF4Q) or 68 kHz (CF4E) and a contact time of 0.4 ms. The CP build-up curves were measured using the π/2 pulse lengths for ^1^H and ^13^C as optimized for frozen and unfrozen samples, and the contact times were varied between 0.1 and 7 ms, with a 1 s recycle delay.

Proton T_2_’ values of the INEPT-detectable carbons were measured at 11.11kHz MAS by inserting a spin-echo delay before the INEPT transfer, with variable delays ranging from 0.18ms to 27ms (multiples of the rotor cycle), and a 1 s recycle delay.

2D-INEPT was performed on CF4E at 40 kHz MAS with a 0.3s recycle delay and the INEPT parameters were chosen for 125 Hz J_CH_ couplings. The ^1^H 90° pulse was set to 3.5 μs, with decoupling at 71 kHz, and the ^13^C 90° pulse was 4.5 μs. The 2D experiment contained 64 points in the indirect dimension and 1200 scans per slice. The CF4E spectrum was compared to CF4Q spectrum collected previously[48] with equivalent parameters except the ^1^H 90° pulse at 3.2 μs with decoupling at 78 kHz.

^13^C chemical shifts were referenced to DSS at 0 ppm using external referencing to adamantane, by setting the downfield peak of adamantane to 40.5 ppm. KBr[52] and a sacrificial aqueous sample with TmDOTP[53] were used to estimate the sample temperatures. At 11.11 kHz, under our decoupling conditions and gas flow, the gas temperature was set to 283K to maintain the sample temperature at 11-13°C for experiments on unfrozen samples, and the gas temperature was set to 247K to maintain the sample temperature at −10 to −15°C for experiments on frozen samples. At 40 kHz MAS, the gas temperature was set to 247K to keep the sample temperature at 23-25°C.

### NMR Data Analysis

Topspin 4.0.7 was used for data processing. 100 Hz of exponential line broadening was used to process the 1D data. Dynamics (Bruker) was used to fit the data points for the T_2_’ measurements. For the 2D data, both dimensions were processed with zero filling to twice the number of data points, sine bell multiplication, and baseline correction (BC_mod = quad). CCPNMR Analysis software was used for peak picking and measuring peak volumes in 2D spectra. The lowest contour level was set to 10 times the noise level in the CF4E spectrum and the same contour levels were set for the CF4Q spectrum used for comparison. Peak volumes were measured using Gaussian fitting integration method.

The Python script used to search for flexible segments to match the NMR peak volume data used the following equation to calculate the score:

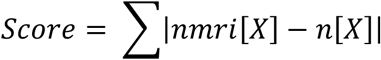

where *nmri[X]* is the NMR intensity for a resonance assigned to amino acid X, and n[X] is the number of times that amino acid is encountered in the stretch of sequence analyzed. The score is the sum of these deviations for all amino acids, so a lower score means a better fit to the NMR data.

The CP-build up curves for both mutants were fit to the equation described in Fig 4 using ProFit 7.

**Figure 1.**
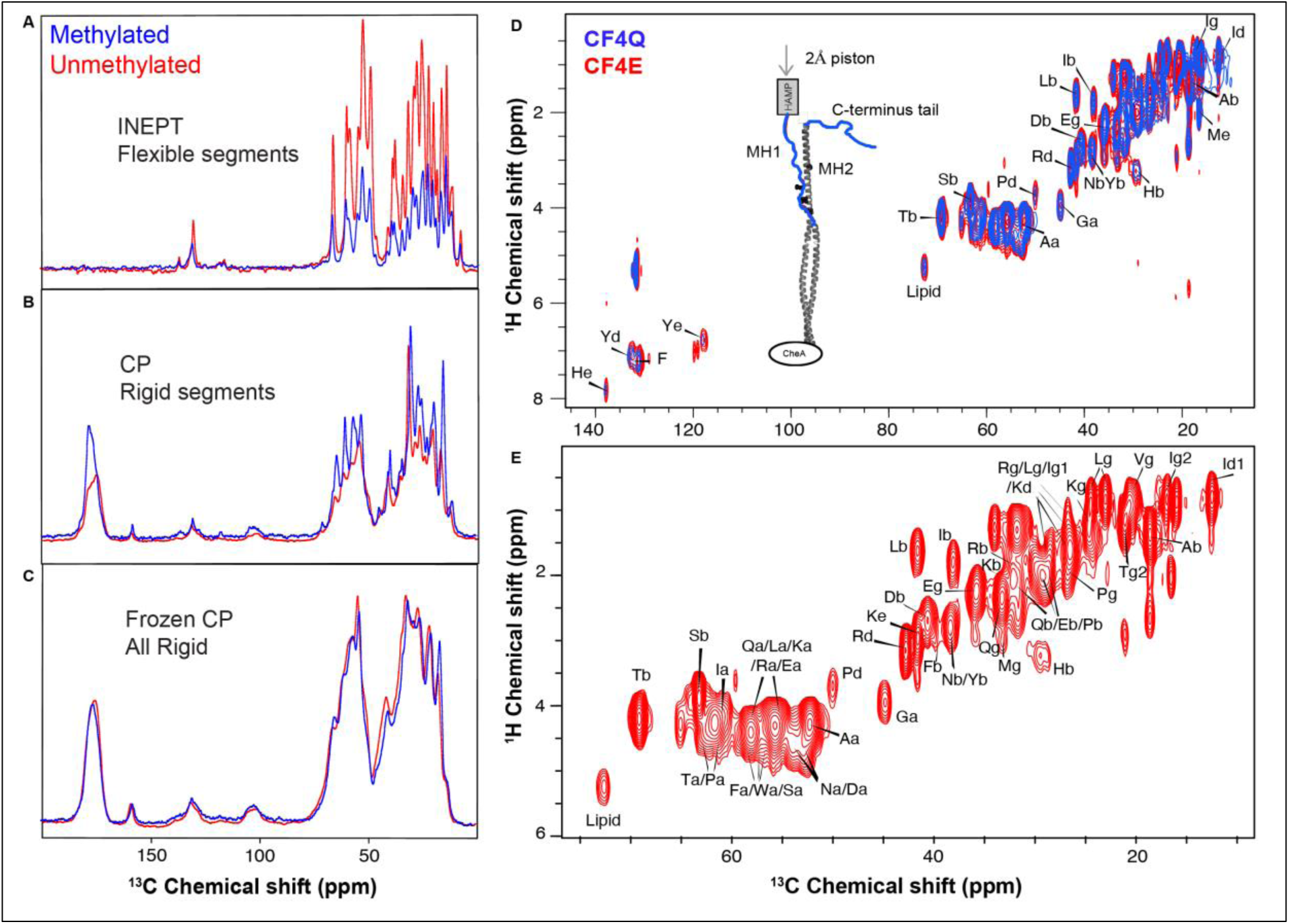
Comparison of NMR spectra of methylated (CF4Q, blue) and unmethylated (CF4E, red) U-^13^C,^15^N-CF in vesicle-mediated complexes with CheA and CheW reveals greater flexibility in the unmethylated receptor. (A) ^13^C INEPT spectra detect flexible segments; 2-fold higher intensity indicates that CF4E is more dynamic. (B) ^13^C CP spectra detect rigid segments; decreased intensity indicates a reduction of rigidity in CF4E complexes. (C) CP intensities increase upon freezing to comparable values for CF4Q and CF4E complexes. (D) ^1^H-^13^C INEPT of U-^13^C,^15^N CF4Q (blue) and U-^13^C,^15^N CF4E (red) in functional complexes show that CF4E has higher intensity than CF4Q. The peaks for CF4Q were previously assigned to the MH1 region and C-terminal tail[48]. The higher intensity of CF4E indicates a longer overall ^1^H T_2_’ and/or additional dynamic segments in the receptor. (E) The CF4E resonances (0-80 ppm region) are identified based on average chemical shifts from BMRB as previously done for CF4Q. All 1D experiments were recorded with 512 scans and a recycle delay of 5 s, at 11.11kHz MAS. 2D ^1^H-^13^C INEPT spectra were collected at 40kHz MAS with 1200 scans, and 0.3s recycle delay. Identical contour levels are shown for both samples. The structural model for CF is based on the 1qu7 crystal structure [Four helical-bundle structure of the cytoplasmic domain of a serine chemotaxis receptor (PDB ID: 1QU7)] [58].

**Figure 2.**
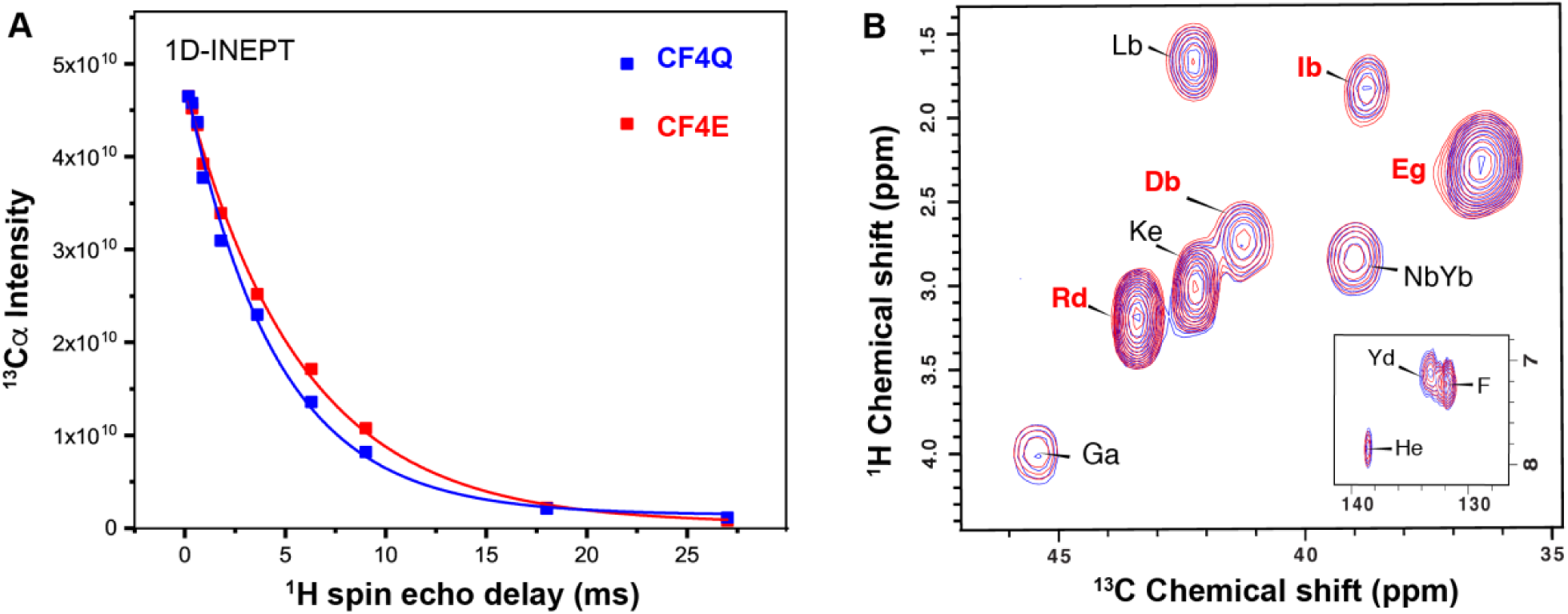
Comparison of proton coherence lifetime (T_2_’) and INEPT peak intensities for methylated (CF4Q, blue) and unmethylated (CF4E, red) U-^13^C,^15^N-CF in vesicle-mediated complexes with CheA and CheW. (A) The ^1^H-T_2_’ of the INEPT-visible (flexible) regions in CF4E is higher than CF4Q sample, accounting for some of the higher overall intensity of the CF4E INEPT spectrum. (B) Representative region of 2D INEPT spectra of CF4Q (blue) and CF4E (red). The CF4E intensity is reduced such that the Tyr peak intensities in both spectra are identical (inset). With this scaling, some residues (black labels) have similar intensities in both samples whereas others (red labels) retain a higher intensity, indicating that this intensity may be coming from an additional dynamic segment in CF4E complexes that is not flexible in CF4Q complexes. Measurements of the ^1^H T_2_’ of the INEPT-visible regions were recorded with 11.11kHz MAS and 1 s recycle delay. Integrals from the Cα region (45-67 ppm) were used to calculate the intensities for T_2_’ estimation; intensities were normalized for an equivalent starting intensity at the shortest delay time.

**Figure 3.**
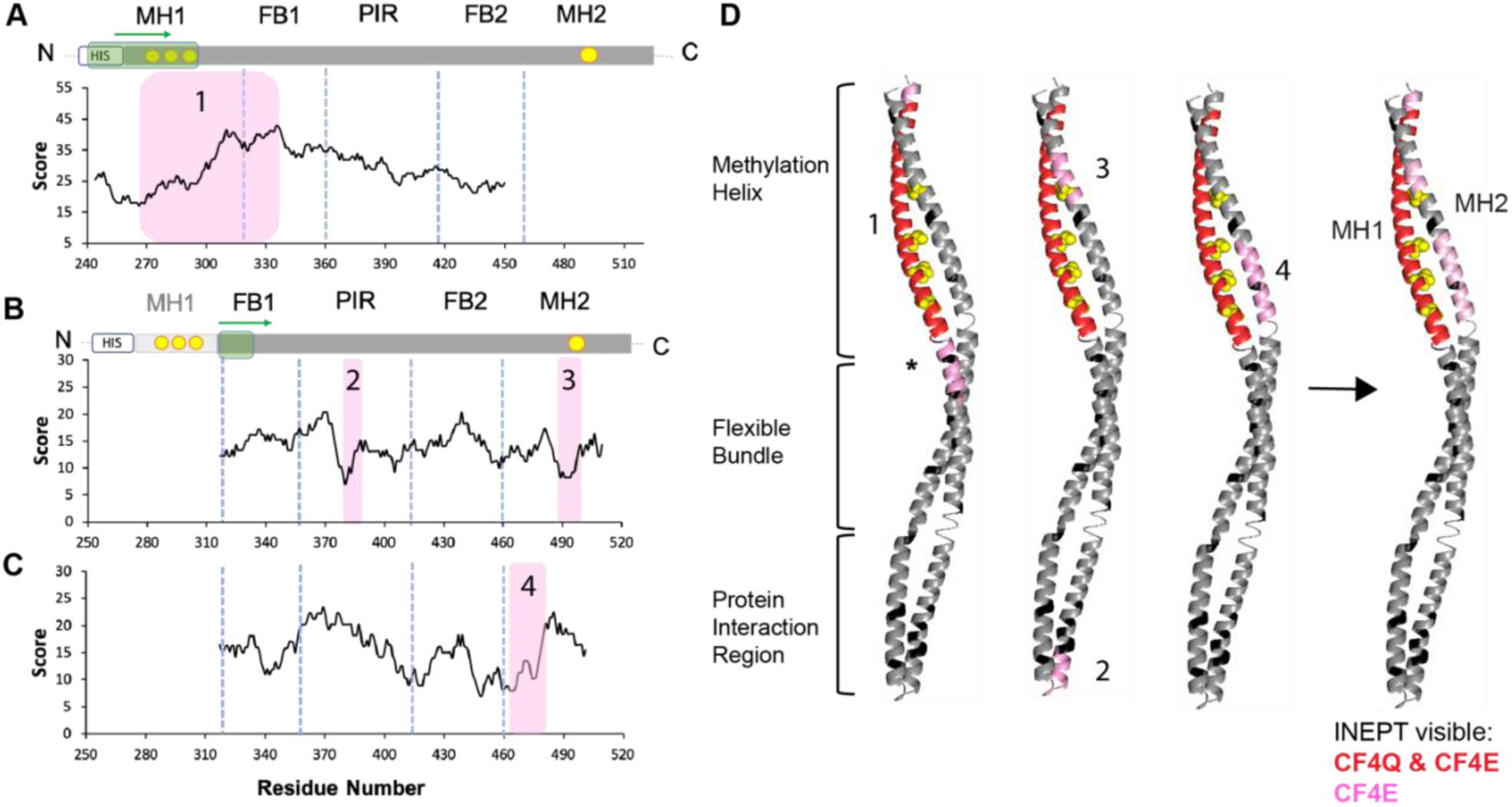
Identification of the likely additional dynamic segment in CF4E. (A) Searching for additional amino acids across the full length CF4E amino acid sequence (without the C-terminal tail) using a single 76-residue sliding window predicts a segment that extends through MH1 and into the flexible bundle (segment 1 highlighted in pink; score of 17. (B) Scanning a 10-residue window along the protein sequence without MH1 and the C-terminal tail identifies two most probable regions in the PIR (segment 2) and MH2 (segment 3) with lowest scores. (C) Scanning a 19-residue window along the same sequence as B identifies an additional segment 4 in MH2. (D) These possible dynamic segments are mapped on the structure of a CF monomer, that includes the locations of the dynamic MH1 previously identified in CF4Q (red), disallowed amino acids that do not show an increase in peak volume in CF4E relative to CF4Q (black), and the methylation sites (space fill yellow). Segment 1 (first structure), identified with the 76-residue scan, contains disallowed residues near the asterisk. The second structure shows segment 2 in the PIR, which is unlikely to be dynamic (see text), and segment 3 in MH2, which contains the fourth methylation site. The third structure shows segment 4 in MH2, which is opposite the MH1 methylation sites. The final structure (right side) represents the likely additional mobile regions in CF4E in pink, based on scoring (A-C) and manual analysis of the sequence. For A-C, the scoring function is described in the text. Scores are plotted for the starting residue of the window, and the lowest scores indicate the least violations of the NMR data. Yellow circles indicate the methylation sites; green arrows represent the direction of the sliding windows (green box).

**Figure 4.**
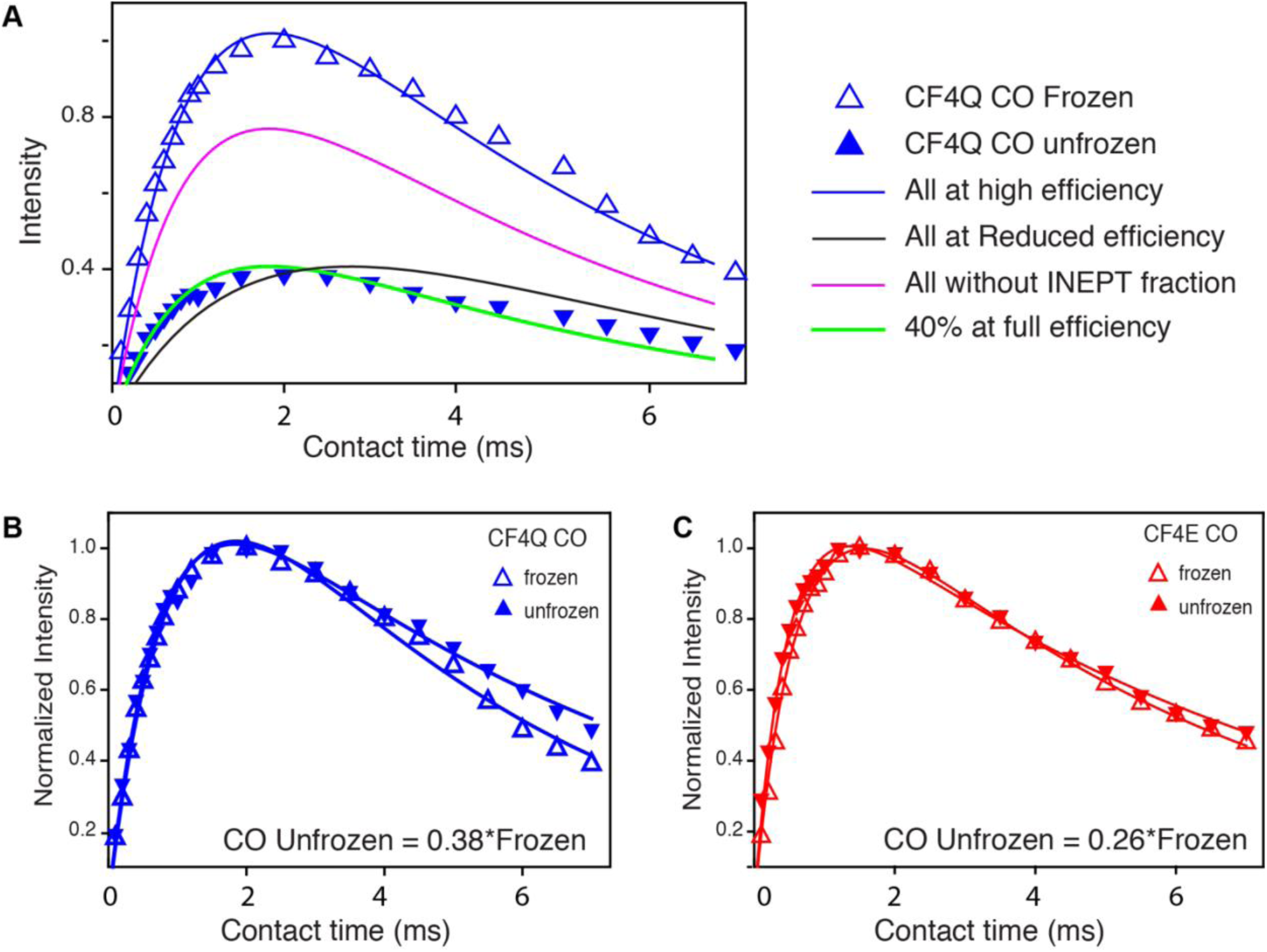
Reduced CP intensity and CP build-up curve shapes suggest CF includes a relatively rigid CP-observable fraction and a CP-invisible fraction. (A) Maximum CP intensity for unfrozen CF4Q (filled triangles) is ∼40% of the frozen CP intensity (open triangles). Blue curve is best fit of frozen CP data to the equation I = M_0_[exp(-t/T_1ρ_) - exp(-t/T_CH_)]/(1+ T_CH_/T_1ρ_), with T_CH_ and T_1ρ_ values given in Table 5. If CP detected all but the INEPT-observable resonances with high efficiency (no motional averaging of the proton-carbon dipolar coupling), the CP build-up should follow the magenta curve (M_0_ scaled to 73%, and no change in T_CH_ and T_1ρ_). Two limiting cases can account for the reduced CP intensity observed for unfrozen CF4Q: (1) If all the residues in the protein exhibit a reduced CP efficiency due to motional averaging of the carbon-proton dipolar coupling, the curve will be shifted towards longer contact times (black curve with M_0_ scaled to 73%, T_CH_ increased 2-fold, and T_1ρ_ unchanged); (2) If only a fraction of the sample is CP-observable (40% for CF4Q) and all observable residues show high CP efficiency, the T_CH_ will be unchanged (green curve with M_0_ scaled to 40%, and both T_CH_ and T_1ρ_ unchanged). The data match case (2), suggesting 40% of CF is relatively rigid (high CP efficiency), and 33% is CP-invisible due to motional averaging of the carbon-proton dipolar coupling. (B) and (C) CP build-up curves for U-^13^C,^15^N CF in functional complexes with CheA and CheW at frozen (open triangles) and unfrozen (filled triangles) temperatures. Integrated intensities of CO (170-190 ppm) are normalized to match the maximum intensity. The similar T_CH_ values (see Table 5) for CF4Q (B) and CF4E (C) indicate that for both states CP observes a relatively rigid fraction of the protein, and the reduction in CP intensity is due to a CP-invisible fraction. The intensities for the frozen sample are higher than those for the unfrozen sample: the CO unfrozen is 0.38*frozen for CF4Q, and 0.26*frozen for CF4E.

## Results

### INEPT-based identification of highly mobile segments: the unmethylated chemotaxis receptor cytoplasmic domain is more dynamic

We have previously used INEPT-based experiments to detect highly mobile segments in the chemoreceptor cytoplasmic fragment (CF4Q) in vesicle-mediated functional complexes with its cytoplasmic partners CheA and CheW[48]. The mobile segments were identified using both deletion of specific regions in the protein and intensities of resolved resonances, especially those of amino acids rare within the protein sequence. These experiments demonstrated that both methylation helix 1 (MH1) and a 34-residue C-terminal tail of CF4Q are highly flexible within functional receptor complexes[48]. Here, we employ similar strategies to compare these dynamics to another mechanistic state of the protein. Comparison of the dynamics of the fully methylated and demethylated receptor should provide insight into the mechanism of receptor adaptation to an ongoing stimulus. The methylation extremes are represented by two mutants, CF4Q and CF4E. In CF4Q, the four methylation sites are mutated to Gln residues that mimic the methylated state of the receptor[54]; CF4E receptor has Glu residues at these sites representing the unmethylated receptor. The NMR experiments are performed on CF bound to vesicles via an N-terminal His tag, and assembled into functional complexes with CheA and CheW[55]. These complexes spontaneously assemble into a hexagonal nanoarray comparable to those observed in bacteria[56] (Figure 7 illustrates one hexagon of the array).

**Table 1:**
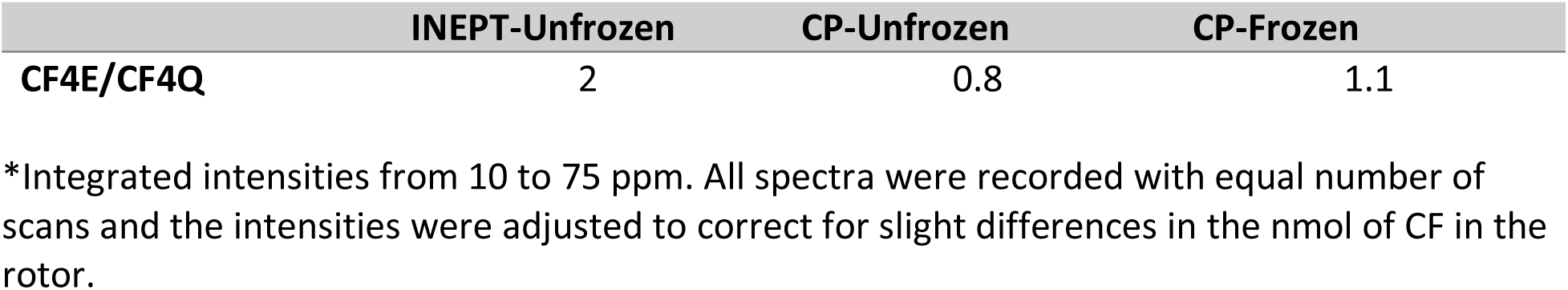
Ratio of CP and INEPT integrated intensities* between the unmethylated (CF4E) to methylated (CF4Q) receptor in vesicle-mediated arrays with CheA and CheW.

**Table 2:** Chemical shifts of resolved amino acids from CF4E and CF4Q in the 2D-^1^H ^13^C INEPT experiments compared to random coil values.

| | Random coil | CF4E | CF4E $\Delta\text{CS}$ | CF4Q | CF4Q $\Delta\text{CS}$ |
| --- | --- | --- | --- | --- | --- |
| **Ala Ca** | 52.1 | 52.95 | **0.85** | 52.87 | **0.77** |
| **Gly Ca** | 45.2 | 45.50 | **0.30** | 45.40 | **0.20** |
| **Ala Cb** | 19.3 | 19.30 | 0.00 | 19.20 | -0.10 |
| **Asp Cb** | 41.2 | 41.29 | 0.09 | 41.21 | 0.01 |
| **His Cb** | 30.1 | 30.15 | 0.05 | 30.06 | -0.04 |
| **Ile Cb** | 38.7 | 38.78 | 0.08 | 38.70 | 0.00 |
| **Leu Cb** | 42.5 | 42.28 | **-0.22** | 42.17 | **-0.33** |
| **Ser Cb** | 64.1 | 63.89 | **-0.21** | 63.86 | **-0.24** |
| **Thr Cb** | 69.7 | 69.83 | 0.13 | 69.80 | 0.10 |

**Table 3:** The ^1^H T_2_’ of the INEPT fraction is higher for ^13^C, ^15^N -CF4E in vesicle-mediated functional arrays with CheA and CheW

| $^1\text{H}$ $T_2'$ of INEPT fraction* | CF4Q | CF4E |
| --- | --- | --- |
| Ca (45-67 ppm) | 4.5 ms | 5.7 ms |
| Full spectrum (10-75 ppm) | 4.48 | 5.8 |
| Thr Cb | 5.58 | 6.68 |
| Ala Cb | 9.35 | 9.35 |
| Glu Cg | 5.4 | 7.4 |

\* $T_2'$ was calculated by fitting the integrated intensities of the indicated regions or peaks to an exponential decay curve.

**Table 4:** Identification of flexible residues based on peak volumes in ^1^H-^13^C-INEPT spectrum of ^13^C,^15^N-CF4E assembled in functional arrays on vesicles with CheA and CheW.*

| Residues | Residues in MH1 | Residues in C-terminus | MH1 + C terminus (flexible in CF4Q) | CF4E peak volumes | Additional flexible residues in CF4E |
| --- | --- | --- | --- | --- | --- |
| Ala $\text{C}\alpha$ | 9 | 3 | 12 | 19.4 | 7.4 |
| Arg $\text{C}\delta$ | 3 | 3 | 6 | 9.2 | 3.2 |
| **Asn $\text{C}\beta$ + Tyr $\text{C}\beta$** | **1+1** | **2** | **4** | **4.2** |  |
| Asp $\text{C}\beta$ | 2 | 1 | 3 | 5.1 | 2.1 |
| Glu $\text{C}\gamma$ | 8 | 3 | 11 | 13.0 | 2.0 |
| Gly $\text{C}\alpha$ | 3 | 0 | 3 | 3.6 | 0.6 |
| **His $\text{C}\epsilon$** | **1** | **0** | **1** | **1.0** |  |
| Ile $\text{C}\beta$ | 2 | 1 | 3 | 4.1 | 1.0 |
| Leu $\text{C}\beta$ | 3 | 2 | 5 | 4.1 | -1.0 |
| Met $\text{C}\epsilon$ | 1 | 0 | 1 | 2.9 | 1.9 |
| **Phe** | **0** | **1** | **1** | **0.8** |  |
| Pro Cδ | 0 | 7 | 7 | 1.2 | |
| Ser Cβ | 5 | 2 | 7 | 12.0 | 5.0 |
| Thr Cβ | 6 | 3 | 9 | 12.0 | 3.0 |
| **Tyr Cε** | **1** | **0** | **1** | **1** |  |
| **Non-Gly Cα** |  |  |  | 106.3 |  |
| Total Cα = Non-Gly + Gly | | | 84 | 109.9 | |

\*Bold rows denote residues with no significant change from CF4Q peak volumes.

**Table 5:** T_CH_ and T_1rho_ calculated from the CP-build curves for CF4Q and CF4E.*

| $T_{CH}$ | CO | | CA | |
| --- | --- | --- | --- | --- |
|  | Frozen | Unfrozen | Frozen | Unfrozen |
| CF4Q | $0.9 \pm 0.004$ | $0.7 \pm 0.003$ | - | - |
| CF4E | $0.6 \pm 0.02$ | $0.5 \pm 0.02$ | - | - |

| $T_{1\rho}$ | CO | | CA | |
| --- | --- | --- | --- | --- |
|  | Frozen | Unfrozen | Frozen | Unfrozen |
| CF4Q | $4.5 \pm 0.2$ | $6.5 \pm 0.3$ | $4.6 \pm 0.05$ | $6.5 \pm 0.07$ |
| CF4E | $5.8 \pm 0.2$ | $6.9 \pm 0.2$ | $4.4 \pm 0.1$ | $5.7 \pm 0.2$ |

\*Integrated intensity for CO (170-190 ppm) and $C\alpha$ (50-75 ppm) were fit to the equation listed in the Figure 4 legend.

Increased intensities in INEPT spectra of CF4E in functional complexes demonstrate greater mobility of the unmethylated receptor. As shown in Fig 1A and Table 1, the INEPT signal for CF4E has 2-fold greater intensity than that of CF4Q. The CP intensity shows a small decrease for CF4E relative to CF4Q (Fig 1B). As expected in the frozen samples, which should be uniformly rigid, the CP intensities for both CF4Q and CF4E are similar (Fig 1C). To gain further insights into the residues that are involved in the increased INEPT signal, we recorded a 2D ^1^H,^13^C refocused INEPT spectrum (Fig 1D). When compared to the 2D ^1^H,^13^C refocused INEPT spectrum we previously reported for CF4Q[48], no new resonances are observed in the CF4E spectrum. However, the overall peak intensity for CF4E is higher than that of CF4Q, indicating a significant increase in dynamics. No significant shifts in the resonances are observed in the CF4E spectrum, indicating the increased dynamics did not cause large changes in the overall structure (Table 2). Comparison of the chemical shifts of both mutants with random coil chemical shifts[57] shows a positive deviation in the Ca values and negative shift in some of the Cb resonances in both mutants, which could be indicative of residual helical structure. This trend was observed for all changes ≥ 0.2 ppm (bold in Table 2). However, these deviations are small, indicating that the dynamic residues identified in the INEPT spectrum have primarily random coil structure. Though not different in structure, the remarkable change in dynamics between the two methylation states is likely to be relevant to the signaling mechanism, and it is important to identify the source of the increased dynamics.

Possible reasons for the increase in INEPT intensity are (1) the ^1^H T_2_’ of the mobile segments increases, and/or (2) other segments (in addition to MH1 and the C-terminal tail) become dynamic enough to be detected in INEPT spectra. We inserted a proton spin echo segment prior to the INEPT transfer to collect a series of 1D INEPT spectra and measure the ^1^H T_2_’ (coherence lifetime) of the INEPT-detectable fraction of both CF4Q and CF4E complexes. We found that indeed the ^1^H T_2_’ of the Cα region was somewhat longer for the unmethylated (CF4E) receptor (Fig 2A and Table 3). For the 4 ms total INEPT delay, the increase in ^1^H T_2_’ from 4.5 to 5.7 ms predicts a 1.2-fold intensity increase in the INEPT signal, accounting for part but not all of the increased INEPT signal. The few resolved resonances in the 1D spectra also showed a similar trend of higher T_2_’ for CF4E residues, with the exception of Ala Cb that showed no change between the two states but had a higher value than other residues (Table 3). This longer T_2_’ for Ala Cb is likely due to the motion of the methyl group that may dominate a smaller change in T_2_’ between CF4Q and CF4E.

Analysis of intensities of resolved peaks in the 2D spectra reveals that some residues show greater intensity increases than others, suggesting the possibility of a newly mobile region in CF4E complexes. To identify this region, we scaled the INEPT spectra to equalize the intensity of the Cd resonance of the single Tyr residue in the CF sequence in INEPT spectra of both the CF4Q and CF4E. This corrects for any changes in intensity due to an overall greater mobility of the Tyr-containing MH1 segment in CF4E. In these scaled spectra we observe that some residues in CF4E remain more intense than in CF4Q (Fig 2B), which may indicate that an additional segment of CF4E has sufficient mobility to be detected in INEPT spectra. As illustrated in Figure 2B, resonances marked with black labels such as Asn/Tyr-Cb and Gly-Ca have equivalent intensities in spectra of CF4Q and CF4E, but those marked with red labels such as Arg Cd and Asp Cb have higher intensity in CF4E.

We employed peak volume analysis of INEPT spectra as previously described[48] to identify the likely additional segment(s) of CF4E with sufficient mobility for INEPT detection. After scaling both CF4Q and CF4E spectra to give peak volumes of two for Tyr Cd, the peak volumes were then divided by the number of unresolved correlations per residue (5 for Phe, 2 for Tyr Cd, and 1 for all other resonances), to give normalized values equal to the number of residues contributing to the peak. This calculation suggests that 110 residues (106 non-Gly Ca + 4 Gly) are observed in INEPT spectra of CF4E (Table 4), which is 26 more than the 84 residues observed in INEPT spectra of CF4Q. The mobile region in CF4Q was previously identified as 34 residues from the C-terminal tail and 50 from methylation helix 1 (MH1)[48]. If the same regions are mobile in CF4E, then the peak volume analysis suggests that an additional region of ∼26 amino acids has similar dynamics. The additional flexible region(s) should include the residues listed in the last column of Table 4, which represent the differences between the peak volumes in the CF4E spectrum and the 84 flexible residues identified in INEPT spectra of CF4Q.

Identification of the likely additional flexible region(s) in CF4E is based on the following criteria. These regions should (1) contain the residue numbers in the last column in Table 4, and (2) not contain the residues listed in bold in the other columns that showed no intensity increases. Also, since the residues listed in the last column sum to ∼26, as expected from the Cα intensities, the additional flexible region(s) should not contain very many other residues. Finally, in order to have sufficient mobility to be INEPT-observable, we suggest that flexible regions contain at least 7 sequential residues (2 turns of a helix). For an unbiased analysis, we used a Python script to identify the most likely additional mobile segment in the sequence that includes the extra NMR intensity observed in INEPT spectra of CF4E. A sliding window of N amino acids was used to scan the CF4E sequence, and compute the difference between the observed NMR intensity for each resolved residue type (‘nmr[X]’ from peak volume analysis) and the number of times that residue occurs in that window ‘n[X]’. The final score for each segment was equal to the sum of the deviations between nmr[X] and n[X] for all of the resolved amino acid resonances, with lower score indicating better fit with the NMR data (see Methods).

The results of this analysis to identify the most likely additional dynamic region in CF4E are presented in Figure 3. The C-terminal tail has been shown to be a flexible tether to the pentapeptide at the C terminus that serves as the binding site for the methylation enzymes[59,60]; thus we excluded the tail residues from our analysis. We used a window of 76 residues (= 110 total - 34 tail) to search the remaining sequence for the best fit to the NMR data and found segment 1, residues 266-330, with the lowest score (shown in pink in Fig 3A; note that the scores are plotted at the starting residue of the window). However, note that the score of 17 reflects a high number of deviations from the NMR data, because this segment includes some disallowed residues (bold in Table 4). This is also shown in Figure 3D, where disallowed residues are represented in black throughout the structure of a CF monomer. Segment 1 is represented in the first structure in Figure 3D, with red representing the 50-residue MH1 region that was previously shown to be flexible in CF4Q[48], and pink representing the additional residues within the 76-residue window. There are no disallowed residues in the N-terminal pink extension, indicating that the flexible MH1 may extend by 3 residues (TVT) in CF4E, accounting for 2 of the 3 flexible Thr residues. However, extension towards the C terminus immediately includes two disallowed Asn residues (**N**AD**N**) indicated by the asterisk in Fig 3D. This analysis confirmed that MH1 is mobile in both CF4E and CF4Q, which is also corroborated by a previous HDX-MS (hydrogen deuterium exchange mass spectrometry) study in the lab showing that the MH1 region shows very rapid HDX in both methylation states[31].

Next, we concentrated on the additional 26 mobile residues in CF4E and omitted the MH1 and C-terminal tail residues from our analysis. Since there was no stretch of the sequence containing 26 amino acids without violations, we used a 10-residue window to scan the rest of the receptor. As shown in Figure 3B, this search identified segments 2 and 3: PIR (protein interaction region, residues 379-389) and MH2 (methylation helix 2, residues 489-498). These segments are highlighted in pink in the second structure of Figure 3D. Since segment 3 in the MH2 region includes a methylation site, it could certainly be more dynamic due to electrostatic repulsion in the demethylated state. Segment 2 is a part of the protein interaction region, which we have previously shown in a hydrogen exchange study to have the slowest deuterium uptake of the whole protein for both CF4Q and CF4E[31]. Thus it is highly unlikely that segment 2 gains sufficient mobility to be INEPT-visible in the unmethylated state of the receptor. Interestingly, both segments 2 and 3 are rich in Ala residues (at least 5 alanines in each), suggesting that the scoring is dominated by the need to fit the largest peak volume increase observed in the INEPT spectra of CF4E (+7 Ala). To account for these Ala, MH2 residues within 491-498 can be definitively assigned to be INEPT-visible in CF4E. For further analysis we removed 5 Ala and 2 Thr residues from the nmr[X] peak volume data and re-ran the script using a 19-residue window. The 2 Thr were removed because these were found in the N terminal extension of MH1 mobility discussed above, but note that the Python analysis identified the same segment whether or not these 2 Thr were included. This search with a 19-residue window yielded the lowest score for segment 4 of MH2 (465-482) (Fig 3C). Although this region has one residue violation (473L), the rest of the amino acids matched the peak volume data set. This identified region is also opposite to the methylation sites on MH1 (Fig 3C). Another segment (448-467) was also identified by the program with similar score as segment 465-482. However, this segment contains 2 Gly, 2 Ile, and 2 Glu residues, which results in 3 overall violations of the NMR data (allowed: 1 Gly, 1 Ile, and 2 Glu, but 1 Glu is already present in segment in 3). By combining this manual data interpretation with the Python scores, we were able to identify the most likely additional highly flexible regions in CF4E are 264-266 (TVT) extending the flexible MH1, and MH2 segments 465-482 (SRGIDQVA**L**AVSEMDRVT) and 491-497 (ESAAAAA). These are shown in the context of the full sequence in Figure S1. This assignment accounts for all of the additional NMR intensity observed in CF4E except for 1 Arg, 1 Met, and 2 Ser, and has only one violation (bold Leu). Physiologically, an increased flexibility of the methylation bundle in the unmethylated receptor will increase the accessibility of the methyltransferase (CheR) to the methylation sites in the receptor. However, further experiments will be required to completely understand the functional relevance of this dynamic difference between the two methylation states of the receptor.

### CP efficiencies and build-up curves suggest that parts of the receptor have intermediate timescale dynamics

Once the INEPT-observable residues were identified, we focused our attention on the remaining less dynamic regions of the receptor. Of the total 310 residues in CF, 84 and 110 were identified as INEPT-visible in CF4Q and CF4E, respectively. If the rest of the protein were rigid on the timescale of the carbon-proton dipolar coupling, we would observe 226 residues for CF4Q (73%) and 200 residues for CF4E (65%) in the CP spectrum. As shown in Figure 4A, comparison of the CP intensity of the frozen CF4Q CO resonance (open triangles), to unfrozen CF4Q at 12°C (filled triangles) demonstrates considerably less signal than expected. In fact, the maximal CP intensity for the unfrozen CO resonance is 38% of that of the frozen sample for CF4Q and 26% for CF4E (based on the scaling needed to superimpose the CP build-up curves shown in Figure 4B-4C). Reduced CP intensity can result from dynamics that partially average the CH dipolar coupling or decrease the ^1^H T_1ρ_. The CP build-up data and curves plotted in Figure 4A are used to deduce whether such dynamics reduce the CP efficiency for CF4Q in functional complexes. The best fit parameters for the CP build-up curves are listed in Table 5. Note the CP build-up is too rapid to determine T_CH_ for the Ca resonance, so Figure 4 focuses on CP build-up for the CO resonance and the other curves are shown in Figure S2. The blue curve in Fig 4A is the best fit for the frozen sample, providing T_CH_ and ^1^H T_1ρ_ values (Table 5) for a relatively rigid protein backbone and efficient CP for all residues. If CP observed all but the INEPT-visible residues with high efficiency (no change in T_CH_ and T_1ρ_), the data would fit to the magenta curve in Fig 4A, which corresponds to a 73% scaling of the blue curve. Instead the data fit to the green curve, corresponding a 40% scaling of the blue curve. This indicates that 40% of the protein backbone has high CP efficiency, comparable to that of frozen samples. The other option is that a longer T_CH_ due to protein dynamics decreases CP efficiency. This is represented by the black curve in Fig 4A, which reaches a comparable intensity at a longer contact time, but does not fit with the observed CP build-up data. The relative rigidity and high CP efficiency of the CP-observable fraction is further demonstrated by the similar values of the T_CH_ and T_1rho_ for frozen and unfrozen CF4Q and CF4E samples, as listed in Table 5 and illustrated by the similarity of the CP build-up curves in Fig 4B and 4C that are scaled to match the maximum intensities. Therefore, both CF4Q and CF4E in functional complexes consist of a relatively rigid fraction with high CP efficiency (comparable to the frozen sample) and a fraction with dynamics on an intermediate timescale, that is neither CP-observable nor INEPT-observable. Based on the frozen/unfrozen CP ratio we calculated that 124 residues of CF4Q and 93 residues of CF4E are CP-observable residues, and thus are rigid on the carbon-proton dipolar coupling timescale (Table 6). Interestingly, the difference in the CP-observable residues between the mutants (−31) is comparable to the difference in the INEPT-observable residues (+26). It remains to be determined whether the additional INEPT-observable mobile region in CF4E complexes is relatively rigid (CP-observable) in CF4Q complexes.

**Table 6.** Estimated number of residues in INEPT and CP spectra of CF4Q and CF4E in vesicle mediated complexes with CheA and CheW*.

|  | Total | INEPT | CP-expected | CP-observed** |
| --- | --- | --- | --- | --- |
| **CF4Q** | 310 | 84 | 226 | 124 |
| **CF4E** | 310 | 110 | 200 | 93 |

\*All calculations are based on the assumption that all CF dimers in the complexes have equivalent dynamics.
\*\*CP-observed residues calculated based on the average unfrozen/frozen CP efficiency observed for the $C\alpha$ and CO for both proteins (CF4Q: CO = 0.38, $C\alpha$ = 0.43, and CF4E: CO = 0.26, $C\alpha$ = 0.32)

## Discussion

### Dynamics and chemoreceptor signaling

We have used solid-state NMR to assess the dynamics on multiple time scales in the *E coli* Asp receptor cytoplasmic fragment (CF) incorporated into vesicle-mediated, functional complexes with CheA and CheW, and demonstrated a difference in dynamics between the two extreme adaptation states of the receptors. Our current understanding of the CF dynamics within these complexes, based on our NMR and HDX-MS studies and consistent with EPR studies discussed below, is summarized in Figure 5, where green represents the rigid protein interaction region of CF that binds CheA and CheW, gray represents regions with intermediate timescale dynamics, and red/pink represent regions with ns-timescale dynamics.

**Figure 5.**
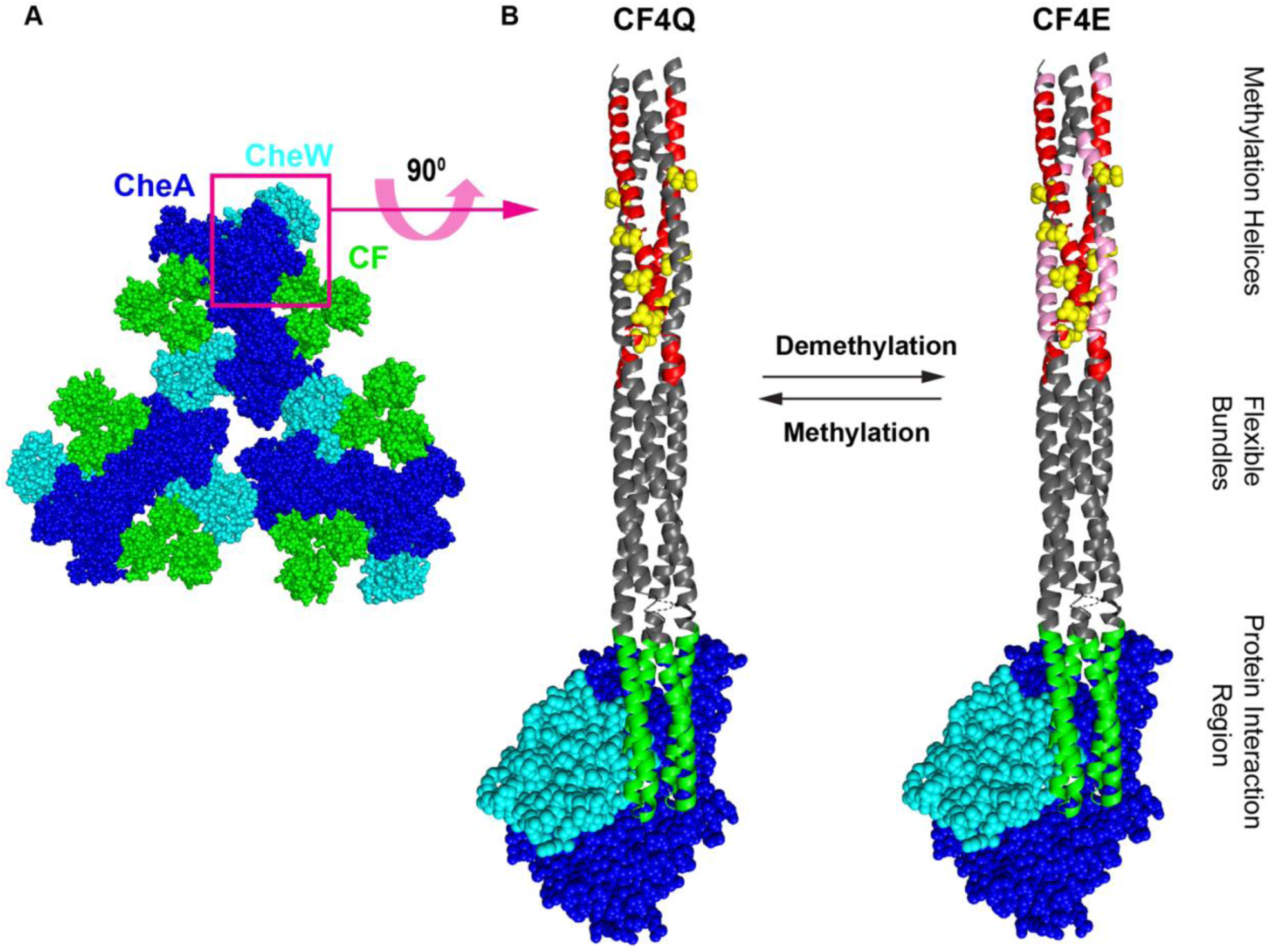
Chemoreceptor dynamics within functional complexes. A. View from below of the array baseplate consisting of the receptor trimer of dimers protein interaction region (green) in complex with CheA dimers (blue) and CheW (cyan). B. View from the side of two methylation states of the complex shown in the red box in A, consisting of a receptor dimer, CheA monomer, and CheW. Receptor dynamics deduced by NMR for these two states are indicated by colors on the receptor dimer. Red and pink indicate INEPT-visible regions with ns or faster timescale dynamics in the methylation region, with red indicating methylation helix 1 (MH1) with these dynamics in both CF4Q and CF4E, and pink indicating additional regions (primarily MH2) with these dynamics in CF4E. Note that the C terminal tail with similar dynamics is not shown. Green indicates the protein interaction region that is likely to be rigid based on NMR observation that only a small fraction of the CF4Q protein has rigid-limit carbon-nitrogen dipolar couplings[47], and hydrogen exchange mass spectrometry evidence that only this region is well-ordered[31]. Regions shown in gray include both CP-observable and CP-invisible segments, with dynamics intermediate between those of the green and red/pink segments. The model is based on docking the 3ur1 crystal structure [TM14 fragment/CheAP4-P5/CheW (PDB ID: 3UR1)] into the electron density observed by cryo-electron microscopy (Structural coordinates courtesy of Brian Crane[36]).

Changes in dynamics of the cytoplasmic domain of the receptor have been proposed to play an important role in signaling[39,40]. Consistent with some of these proposals, EPR studies on fusion proteins indicate that dynamics of both the methylation and the protein interaction regions increase in the 4E demethylated state, which is typically kinase-OFF, relative to the 4Q mimic of the methylated state[61]. Another EPR study corroborated this observation that the methylation region is more dynamic in 4E than in 4Q, and also showed that methylation helix 1 (MH1) is more dynamic than methylation helix 2 (MH2) in both states[62].These EPR studies were done on receptors in the absence of their cytoplasmic partners CheA and CheW. Our previous INEPT NMR study of functional complexes of CF4Q with CheA and CheW[48] demonstrated that the dynamics of MH1 (red region in Fig 5) in CF4Q are comparable to those of the disordered C-terminal tail. Such a flexible connection to the cytoplasmic domain eliminates some possible signaling mechanisms, such as propagation of a conformational change down the entire length of the receptor. Instead, based on HDX-MS measurements, we have recently proposed that signaling occurs via modulation of the partially disordered nature of much of the cytoplasmic domain[31]. Such a mechanism is consistent with observation of more rapid hydrogen exchange throughout CF in both a kinase-off mutant (CF4Q.A411V) and in CF4E[31], as well as increased INEPT intensities for MH1 in both CF4Q.A411V[48] and CF4E (this study). Our current study also shows that the increased dynamics of CF4E (pink segments in Fig 5) include ns timescale dynamics in parts of MH2. This observation is consistent with the EPR results of Bartelli et al[62], who reported mobility parameters for MH2 in CF4E that are in the same range as those of MH1 in both CF4Q and CF4E. The increased dynamics of MH2 in CF4E is likely part of the increased disorder throughout the CF that we propose turns off the kinase CheA. Furthermore, we hypothesize that the increased flexibility of the methylation bundle in CF4E may increase the accessibility of the methylation sites to the methyltransferase when the receptor is in the unmethylated state. Such a change could account for an important feature of chemoreceptor adaption: the kinase-OFF state exhibits a faster rate of methylation.

It is interesting to note that both hydrogen exchange and INEPT NMR data suggest a larger increase in dynamics in the unmethylated CF4E than in the kinase-OFF CF4Q.A411V mutant, relative to CF4Q. Regions with faster hydrogen exchange are more widespread in CF4E than in CF4Q.A411V[31]. INEPT intensities increase only ∼1.3-fold in CF4Q.A411V [48] vs ∼2-fold in CF4E (this study). This may be due to the inequivalence of the CF dimers within the hexagonal array. It is likely that the A411V mutation alters interactions between CF and CheA, and only one of the dimers in each receptor trimer of dimers interacts with CheA. In contrast, the 4Q to 4E change alters properties of all dimers in the CF array. This inequivalence of receptor dimers raises the possibility that dimers exhibit different dynamic properties and these do not change uniformly with signaling state: for example, the receptor trimer of dimers could have a single dimer with an INEPT-visible MH1 in CF4Q, and the INEPT intensity could increase two-fold in CF4E because two dimers have INEPT-visible MH1. Consistent with such possibilities, recent cryo-EM evidence indicates that receptor dimers differ in their structural rigidity in the methylation region[63]. Thus, further studies are needed to investigate whether the three inequivalent receptor dimers in functional complexes have similar dynamic regions and properties.

### Dynamics overview provides insight for NMR experimental design

Analysis of the CP build-up curves for CF4Q and CF4E complexes indicates that a fraction of the receptor in both states shows intermediate dynamics and is invisible to both INEPT and CP NMR experiments performed in this study. Table 6 summarizes the distribution of dynamics in the these CF methylation states: 27-35% highly flexible (INEPT-visible, with ns timescale dynamics), 33-35% with intermediate timescale dynamics (not observed in INEPT or CP spectra), and 40-30% relatively rigid (CP-visible). The small fraction of relatively rigid CF (40—30% CP-observable) is consistent with the proposed partial disorder of the CF within functional complexes [31], and presents both a challenge and an opportunity for NMR studies.

NMR and HDX-MS experiments are synergistic for the identification of dynamic regions of the CF. The INEPT-observable MH1 and C-terminal tail both exhibit complete hydrogen exchange within 3 minutes in both CF4Q and CF4E, consistent with being flexible, unstructured regions. In addition, the 455-472 segment of MH2 exhibits this very fast exchange in CF4E. This overlaps both of the lowest scoring segment identified by the Python analysis of Fig 4C: 448-467 and 465-482. As discussed above, the sequence of the latter segment better fits the NMR data and thus is more likely to be the INEPT-observable MH2 region.

Our previous study of CF4Q complexes identified ms-timescale dynamics within the receptor by observing that 40% of the CP-observable region shows back-bone dynamics that fully average the ∼1 kHz carbon-nitrogen dipolar coupling[48]. Since we have now shown that all CP-observable residues (124 in CF4Q) exhibit full CP efficiency, the number of residues with rigid-limit carbon-nitrogen dipolar coupling can be calculated as 124*0.6 ≈ 75 residues. This rigid core thus accounts for ≈25% of the protein. Since our HDX-MS studies on the receptor in functional complexes with CheA and CheW have shown that only the protein interaction region (57 residues, green in Fig. 5) exhibits hydrogen exchange properties consistent with a fully structured region [31], this region is likely to have rigid-limit carbon-nitrogen dipolar couplings. The extensive partial disorder of the CF could account for the broad line widths we previously observed in NcaCX spectra of frozen samples [64]. Our current knowledge of the CF dynamics predict that NCA/NCO spectra of unfrozen samples will selectively detect only the 75 rigid residues, including the well-structured protein interaction region, which should be amenable to sequence assignments to study the effects of changes in signaling states on receptor-kinase interactions. Thus, NMR and HDX-MS measurements of dynamics of the CF are enabling the design of NMR strategies to tackle a complex system with extensive dynamics and partially disordered regions important in the signaling mechanism of this receptor. These strategies can be easily applied to other protein complexes with disordered regions and dynamics encompassing multiple timescales to enable the study of mechanisms that are fundamental to many cellular processes.

## Acknowledgements

The NMR experiments were collected at the University of Massachusetts Amherst NMR facility, with support from the Institute for Applied Life Sciences. We are grateful to Weiguo Hu and Jasna Fejzo for support for the NMR experiments and data analysis, and to Brian Crane for sharing coordinates for the structural model. This research was supported by National Institutes of Health Grant R01-GM120195. KAW was partially supported by National Research Service Award T32 GM008515 from the National Institutes of Health, as part of the UMass Chemistry-Biology Interface Training Program.

