## Supplementary Material for "Strategies for identifying dynamic regions in protein complexes: flexibility changes accompany methylation in chemotaxis receptor signaling states"

for

**N**- His Tag

LTNKPQTPSRPASEQPPAQPRLRIAEQDPNWETF- **COOH**

| MH1 | | MH2 | |
|---|---|---|---|
| 259 | R | 515 | P |
| 260 | S | 515 | S |
| 261 | L | 515 | A |
| 262 | T | 515 | A |
| 263 | D | 515 | L |
| 264 | T | 514 | R |
| 265 | V | 513 | F |
| 266 | T | 512 | A |
| 267 | H | 511 | S |
| 268 | V | 510 | V |
| 269 | R | 509 | A |
| 270 | E | 508 | Q |
| 271 | G | 507 | T |
| 272 | S | 506 | L |
| 273 | D | 505 | R |
| 274 | A | 504 | S |
| 275 | I | 503 | A |
| 276 | Y | 502 | Q |
| 277 | A | 501 | E |
| 278 | G | 500 | E |
| 279 | T | 499 | L |
| 280 | R | 498 | A |
| 281 | E | 497 | A |
| 282 | I | 496 | A |
| 283 | A | 495 | A |
| 284 | A | 494 | A |
| 285 | G | 493 | A |
| 286 | N | 492 | S |
| 287 | T | 491 | E |
| 288 | D | 490 | Q |
| 289 | L | 489 | V |
| 290 | S | 488 | L |
| 291 | S | 487 | S |
| 292 | R | 486 | A |
| 293 | T | 485 | N |
| 294 | E | 484 | Q |
| 295 | Q | 483 | Q |
| 296 | Q | 482 | T |
| 297 | A | 481 | V |
| 298 | S | 480 | R |
| 299 | A | 479 | D |
| 300 | L | 478 | M |
| 301 | E | 477 | E |
| 302 | E | 476 | S |
| 303 | T | 475 | V |
| 304 | A | 474 | A |
| 305 | A | 473 | L |
| 306 | S | 472 | A |
| 307 | M | 471 | V |
| 308 | E | 470 | Q |
| 309 | Q | 469 | D |
| 310 | L | 468 | I |
| 311 | T | 467 | G |
| 312 | A | 466 | R |
| 313 | T | 465 | S |
| 314 | V | 464 | Q |
| 315 | K | 463 | E |
| 316 | Q | 462 | D |
| 317 | N | 461 | S |
| 318 | A | 460 | A |

| FB1 | | FB2 | |
|---|---|---|---|
| 319 | D | 459 | S |
| 320 | N | 458 | A |
| 321 | A | 457 | I |
| 322 | R | 456 | E |
| 323 | Q | 455 | G |
| 324 | A | 454 | M |
| 325 | S | 453 | I |
| 326 | Q | 452 | D |
| 327 | L | 451 | T |
| 328 | A | 450 | V |
| 329 | Q | 449 | R |
| 330 | S | 448 | T |
| 331 | A | 447 | V |
| 332 | S | 446 | A |
| 333 | D | 445 | N |
| 334 | T | 444 | V |
| 335 | A | 443 | I |
| 336 | Q | 442 | N |
| 337 | H | 441 | N |
| 338 | G | 440 | M |
| 339 | G | 439 | T |
| 340 | K | 438 | E |
| 341 | V | 437 | G |
| 342 | V | 436 | A |
| 343 | D | 435 | S |
| 344 | G | 434 | E |
| 345 | V | 433 | V |
| 346 | V | 432 | L |
| 347 | K | 431 | V |
| 348 | T | 430 | S |
| 349 | M | 429 | G |
| 350 | H | 428 | T |
| 351 | E | 427 | D |
| 352 | I | 426 | V |
| 353 | A | 425 | R |
| 354 | D | 424 | S |
| 355 | S | 423 | V |
| 356 | S | 422 | S |
| 357 | K | 421 | D |
| 358 | K | 420 | E |
| 359 | I | 419 | I |
| 360 | A | 418 | L |

| PIR | | | |
|---|---|---|---|
| 361 | D | 417 | A |
| 362 | I | 416 | K |
| 363 | I | 415 | I |
| 364 | S | 414 | E |
| 365 | V | 413 | K |
| 366 | I | 412 | A |
| 367 | D | 411 | A |
| 368 | G | 410 | Q |
| 369 | I | 409 | A |
| 370 | A | 408 | S |
| 371 | F | 407 | R |
| 372 | Q | 406 | S |
| 373 | T | 405 | A |
| 374 | N | 404 | L |
| 375 | I | 403 | N |
| 376 | L | 402 | R |
| 377 | A | 401 | V |
| 378 | L | 400 | E |
| 379 | N | 399 | G |
| 380 | A | 398 | A |
| 381 | A | 397 | V |
| 382 | V | 396 | V |
| 383 | E | 395 | A |
| 384 | A | 394 | F |
| 385 | A | 393 | G |
| 386 | R | 392 | R |
| 387 | A | 391 | G |
| 388 | G | 390 | Q |
| 389 | E | | |

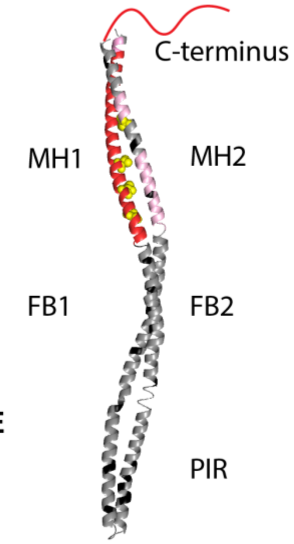


**Flexible in CF4Q and CF4E**
**Flexible in CF4E only**
**Disallowed amino acids**
**Methylation sites**

Fig S1: INEPT-visible residues in both CF4Q and CF4E (red) and additional residues flexible in CF4E (pink) are highlighted on the cytoplasmic fragment sequence. Each table represents adjacent residues in different regions[65] of the hairpin structure: methylation helices (MH1 and MH2), flexible bundles (FB1 and FB2), and the protein interaction region (PIR), with the C-terminal tail at the top of MH2 region. Black arrows denote the directionality of the sequence with respect to the structure. Note that the additional flexible segments in CF4E are near or opposite the methylation sites (highlighted in yellow). The residues that did not show increased peak volume are highlighted in grey. The helix is color coded similar to the tables to depict the regions in the table in context of hairpin structure of the receptor.

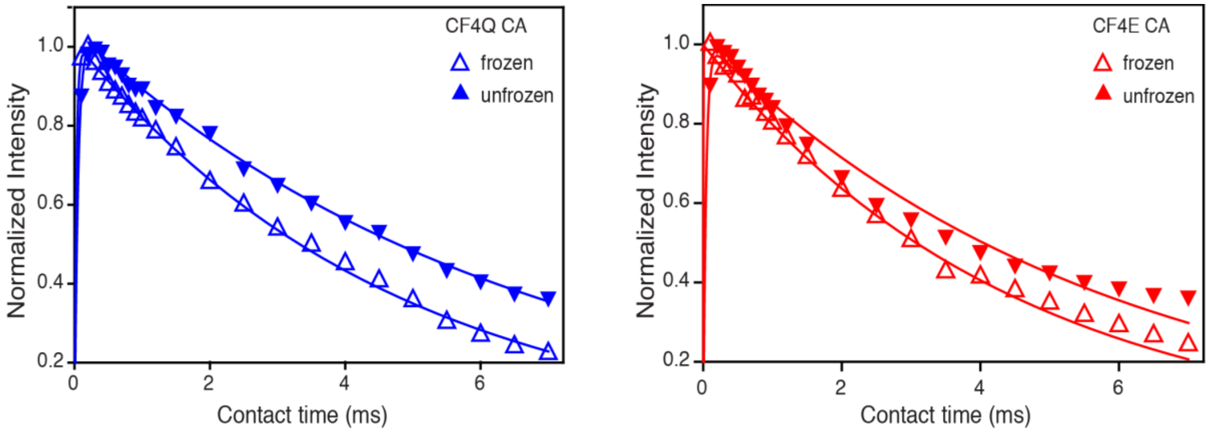


Fig S2: CP build-up curves for U-$^{13}$C,$^{15}$N CF in functional complexes with CheA and CheW at frozen (open triangles) and unfrozen (filled triangles) temperatures. Integrated intensities of Ca (45-70 ppm) are normalized to match the maximum intensity. The rapid build-up for both CF4Q and CF4E did not allow for the precise calculation of $T_{CH.}$ The intensities for the frozen sample are higher than those for the unfrozen sample: the C$\alpha$ unfrozen is 0.43*frozen for CF4Q, and 0.32*frozen for CF4E.
